# Constraining models of dominance for nonsynonymous mutations in the human genome

**DOI:** 10.1101/2024.02.25.582010

**Authors:** Christopher C. Kyriazis, Kirk E. Lohmueller

**Affiliations:** Department of Ecology and Evolutionary Biology, University of California, Los Angeles, USA; Interdepartmental Program in Bioinformatics, University of California, Los Angeles, USA; Department of Human Genetics, David Geffen School of Medicine, Los Angeles, USA

## Abstract

Dominance is a fundamental parameter in genetics, determining the dynamics of natural selection on deleterious and beneficial mutations, the patterns of genetic variation in natural populations, and the severity of inbreeding depression in a population. Despite this importance, dominance parameters remain poorly known, particularly in humans or other non-model organisms. A key reason for this lack of information about dominance is that it is extremely challenging to disentangle the selection coefficient (*s*) of a mutation from its dominance coefficient (*h*). Here, we explore dominance and selection parameters in humans by fitting models to the site frequency spectrum (SFS) for nonsynonymous mutations. When assuming a single dominance coefficient for all nonsynonymous mutations, we find that numerous *h* values can fit the data, so long as *h* is greater than ∼0.15. Moreover, we also observe that theoretically-predicted models with a negative relationship between *h* and *s* can also fit the data well, including models with *h*=0.05 for strongly deleterious mutations. Finally, we use our estimated dominance and selection parameters to inform simulations revisiting the question of whether the out-of-Africa bottleneck has led to differences in genetic load between African and non-African human populations. These simulations suggest that the relative burden of genetic load in non-African populations depends on the dominance model assumed, with slight increases for more weakly recessive models and slight decreases shown for more strongly recessive models. Moreover, these results also demonstrate that models of partially recessive nonsynonymous mutations can explain the observed severity of inbreeding depression in humans, bridging the gap between molecular population genetics and direct measures of fitness in humans. Our work represents a comprehensive assessment of dominance and deleterious variation in humans, with implications for parameterizing models of deleterious variation in humans and other mammalian species.

**Author Summary:** The dominance coefficient (*h*) of a mutation determines its impact on organismal fitness when heterozygous. For instance, fully recessive mutations (*h*=0) have no effects on fitness when heterozygous whereas additive mutations (*h*=0.5) have an effect that is intermediate to the two heterozygous mutations. The extent to which deleterious mutations may be recessive, additive, or dominant is a key area of study in evolutionary genetics. However, dominance parameters remain poorly known in humans and most other organisms due to a variety of technical challenges. In this study, we aim to constrain the possible set of dominance and selection parameters for amino acid changing mutations in humans. We find that a wide range of models are possible, including models with a theoretically-predicted relationship between *h* and *s*. We then use a range of plausible selection and dominance models to explore how deleterious variation may have been shaped by the out-of-Africa bottleneck in humans. Our results highlight the subtle influence of dominance on patterns of genetic load in humans and demonstrate that models of partially recessive mutations at amino-acid-changing sites can explain the observed effects of inbreeding on mortality in humans.

## Introduction

Dominance is a key concept in genetics, determining the fitness effect of a heterozygous genotype compared to that of the two homozygous genotypes. When fully recessive (*h*=0), a mutation has no impact on fitness in the heterozygous state, and when additive (*h*=0.5), the fitness effect of a heterozygote is exactly intermediate to the two homozygous genotypes. The extent to which mutations have additive or recessive impacts on fitness is critical for many aspects of evolutionary genetics. For instance, dominance is a central parameter determining the impact of population size on deleterious variation and inbreeding depression [1–5]. Numerous studies have demonstrated that the behavior of deleterious mutations in a population greatly differs when mutations are highly recessive (*h*<0.05) compared to when mutations are partially recessive (*h*>0.1) [1–5]. Thus, estimating dominance parameters is an essential component of modelling the effects of demography on recessive deleterious variation.

The importance of dominance in human evolutionary genetics has previously been highlighted by numerous studies examining the effects of the out-of-Africa bottleneck in humans on the relative burden of deleterious variation in African and non-African populations [5–10].

Specifically, these studies aimed to determine whether non-African populations may have an elevated burden of deleterious variation (also known as “genetic load”) due to a prolonged population bottleneck that occurred when humans migrated from Africa. A general conclusion that emerged from these studies is that, when deleterious mutations are additive, recent demography appears to have a slight influence on genetic load, whereas under a recessive model, recent demography may have a more pronounced effect on genetic load [5–9]. These findings have important implications for our understanding of how human evolutionary history has impacted the efficacy of natural selection across human populations.

Despite this significance of dominance in evolutionary genetics, dominance parameters remain very poorly quantified in humans and other vertebrates. In humans, almost no estimates of dominance parameters exist, and those that are available are for small subsets of the genome. For instance, Balick et al. [5] devised an elegant approach of contrasting patterns of genetic variation in bottlenecked vs. non-bottlenecked populations, based on the knowledge that bottlenecks impact genetic variation in differing ways when mutations are additive or recessive [4]. Their analysis estimated that genes associated with known autosomal recessive diseases have an average dominance coefficient of *h*=0.2. However, the degree to which this estimate applies to a broader set of deleterious mutations remains unclear. In the relative absence of information about dominance in humans or vertebrates, many studies instead opt to use *ad hoc* dominance parameters for modelling deleterious variation (e.g., [2,6,11]), or explore only the extreme cases of additive and fully recessive mutations (e.g., [9,12]). Thus, our understanding of how demography influences the behavior of deleterious mutations in humans and other species remains greatly limited by a poor understanding of dominance parameters.

In a laboratory setting, several experimental studies have been conducted aiming to quantify dominance using model organisms such as *Drosophila melanogaster* and *Saccharomyces cerevisiae*. These studies have generally found support for deleterious mutations being partially recessive, with mean *h* estimates ranging from ∼0.1-0.4 [13–18]. Moreover, some studies have also found support for a relationship between *h* and *s* (hereafter, *h-s* relationship), where more deleterious mutations tend to be more recessive [14,17,18]. This negative relationship between *h* and *s* was predicted by theoretical models of dominance proposed by Wright and Haldane [19,20]. However, there many caveats associated with these experimental studies. First, experimental manipulations are time-consuming and only a handful of such experiments have been conducted. Consequently, several studies have reanalyzed existing experimental data and found that estimates of *h* may depend greatly on the analytical approach and that such estimates are typically associated with a fair amount of uncertainty [14,15]. Moreover, given that these experiments have been conducted on model organisms in controlled laboratory settings, it remains unclear whether these results are relevant for natural populations in vertebrate taxa such as humans.

Another common approach for estimating selection and dominance parameters relies on using evolutionary models to infer parameters from patterns of genetic variation, as summarized by the site frequency spectrum (SFS; reviewed in [21]). Based on the Poisson random field model [22], Williamson et al. [23] developed an approach to infer dominance using diffusion theory to model the change in allele frequency over time due to genetic drift and selection. Parameters, including *h*, are estimated by finding the values that yield an SFS that is close to that from the empirical data. While this approach is theoretically elegant, it has not been applied to many species. One reason for this is that, when the distribution of fitness effects for new mutations (DFE) is unknown, it is hard to distinguish between different combinations of *s* and *h* [24,25]. Intuitively, most deleterious mutations in natural populations are segregating in the heterozygous state. This provides information about *hs*, rather than *s* and *h* separately [24,25]. Given these challenges, studies that have attempted to estimate the DFE typically ignore dominance entirely by assuming that all mutations are additive (e.g., [26–29]). However, in humans, one previous study attempted to fit non-additive models to the SFS, assuming the same value of *h* for all mutations [30]. This study found that a range of *h* values fit the data reasonably well, so long as *h*>0.3 [30]. However, this study did not attempt to fit an *h-s* relationship, given the challenges associated with identifiability of *h* and *s* parameters.

Previously, Huber et al. [25] circumvented this identifiability issue by leveraging selfing and outcrossing *Arabidopsis* populations for estimating dominance parameters. Genetic variation data from the outcrossing population provided information about *hs* while the selfing population provided information about *s*, because all mutations were found in the homozygous state. Using this framework, Huber et al. [25] found statistical support for an *h-s* relationship for amino acid changing mutations in *Arabidopsis*, with more deleterious mutations being more recessive. Specifically, they found that even moderately deleterious mutations (1e-3<|*s*|<=1e-2) were highly recessive (*h*<0.05), whereas more neutral mutations may be dominant (*h*=1) or additive (*h*=0.5) [25]. These results contrast with previous experimental work, where only very strongly deleterious mutations (|*s*|>0.1) are typically found to be highly recessive (*h*<0.05) [13–18]. Moreover, these results also contrast with previous SFS-based studies in humans, where highly recessive models were shown to have a poor fit to the data [30]. The extent to which these contrasting results may be due to methodological reasons or true differences in dominance parameters across species remains unclear.

Here, we explore the fit of selection and dominance models to patterns of genomic variation in humans. Given that we cannot reliably separate *s* and *h* for based on genetic variation data, we instead constrain the range of dominance models are consistent with the nonsynonymous SFS. We then use these results to parameterize simulations revisiting the question of whether the out-of-Africa bottleneck has led to differences in genetic load between African and non-African human populations. Altogether, our analysis represents a comprehensive exploration of dominance in humans, with numerous implications for understanding the relevance of recessive deleterious variation in humans and other species.

## Results

### Inference of dominance assuming a single dominance coefficient

We first tested whether models with a single dominance coefficient (*h*) can fit the SFS for nonsynonymous mutations in the human genome. To do this, we followed the same approach as Kim et al. [27]. We generated the SFS for synonymous and nonsynonymous mutations from 432 individuals with European ancestry from the 1000 Genomes Project [31], then used the synonymous SFS to infer a demographic model consisting of a bottleneck followed by recent exponential growth (see **Methods** and **Tables S1-S2**). We conditioned on this demographic model for all subsequent inferences.

We began by assuming that the DFE follows a gamma distribution, since previous work has suggested that a gamma distribution is a reasonable functional form for the DFE [26–29]. When assuming that all mutations are additive (*h*=0.5), the best-fitting DFE has a log-likelihood (LL) of - 1450.58 (**Fig. 1a; Table S3**) and the shape (𝛼) and scale (𝛽) parameters inferred assuming *h*=0.5 are similar to those inferred from previous studies for humans [26,27,32]. We then re-inferred the parameters of the gamma distribution for the DFE when assuming different values of *h* (**Fig. 1; Table S3**). We found that the log-likelihood greatly decreased for models with *h*<0.15, suggesting that highly recessive models do not fit the data well (**Fig. S1**), in agreement with previous work [30]. However, other models where all nonsynonymous mutations are partially recessive (*h*=0.35, LL=- 1450.78) or dominant (*h*=0.75, LL=-1450.83) are within 1.92 LL units of the additive model, suggesting that these models also provide a reasonable fit to the data. As the assumed value of *h* become more recessive, the shape parameter of the DFE tends to decrease, while the scale increases (**Table S3**). Consequently, the average selection coefficient (E[*s*]) becomes more deleterious as *h* decreases (**Table S3**), as expected given that the overall product of *h*s* should remain constant.

**Figure 1:**
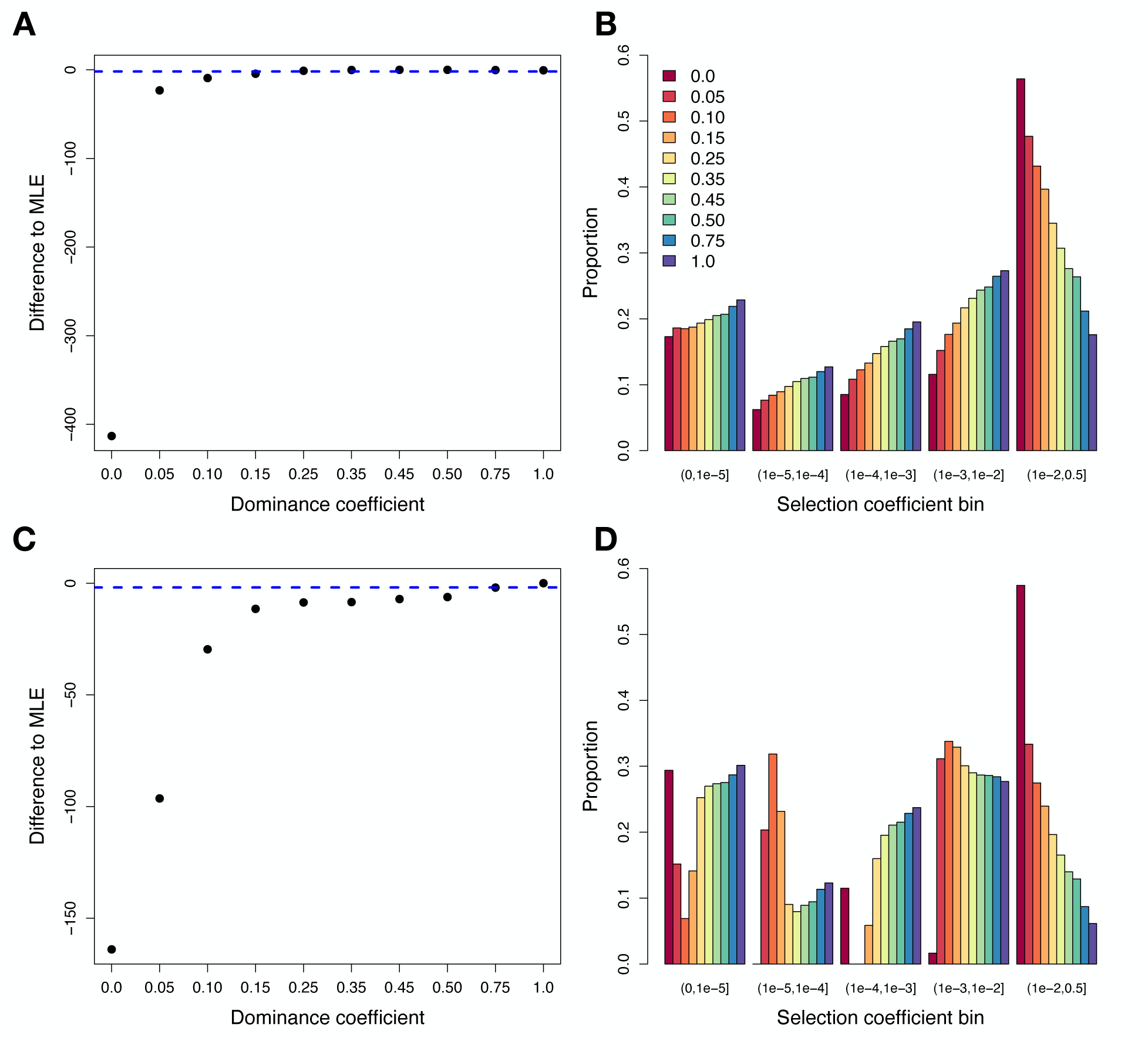
Results when fitting a gamma (top panels) and discrete (bottom panels) DFE model under different dominance coefficients. (A) Profile log-likelihood for the gamma model. The Y-axis shows the difference in log-likelihood relative to the best-fitting model (*h*=0.50, LL=-1450.58) and the dashed blue line depicts 1.92 log-likelihood units below the top model. (B) The proportion of mutations for each selection coefficient bin under the gamma model when assuming different dominance coefficients. Note that the gamma model assumes a continuous distribution but results are shown here as discrete bins to facilitate visualization. (C) Profile log-likelihood for the discrete DFE model. The Y-axis shows the difference in log-likelihood relative to the best-fitting model (*h*=1.0, LL=-1446.76). (D) The proportion of mutations for each selection coefficient bin under the discrete DFE model when assuming different dominance coefficients.

Next, we examined the fit of a discrete DFE including five bins of *s* for new mutations: neutral (0 < |𝑠| ≤ 10^!"^), nearly neutral (10^!"^ < |𝑠| ≤ 10^!#^), weakly deleterious (10^!#^ < |𝑠| ≤ 10^!$^) moderately deleterious (10^!$^ < |𝑠| ≤ 10^!%^), and strongly deleterious (10^!%^ < |𝑠| ≤ 0.5). The discrete DFE quantifies the proportions of mutations in each bin. The advantage of using a discrete DFE is that it does not assume a unimodal distribution for *s*. Further, as shown below, this DFE more easily allows for mutations with different values of *s* to have different values of *h.* When assuming that all mutations are additive, we found that the fit of the discrete DFE to the SFS is slightly worse than the fit of the gamma DFE (LL=-1452.97, 𝛥AIC to gamma=8.8; **Table S4**), consistent with what was observed previously [27]. We then re-inferred the parameters of the discrete DFE when assuming different values of *h* (**Fig. 1C-D**). Under the discrete DFE, we found that models with *h* ranging from 0.15 to 1.0 fit the data well, with a maximum log-likelihood of -1446.76 when *h*=1.0 (**Fig. 1C and S2; Table S4**). However, we observed a very poor fit for models where mutations were highly recessive (*h*<0.15)(**Fig. 1C-D; Fig. S2**). In these highly recessive models, we again observe that the proportion of mutations that are strongly deleterious increases substantially (**Fig.1D**), with nearly 60% of new mutations inferred to have |𝑠| > 10^−2^ when *h*=0.0. By contrast, when *h*=1.0, only ∼6% of mutations are inferred to have |𝑠| > 10^!%^(**Fig. 1D**).

### Fitting models with multiple dominance coefficients

Our analyses above assumed a single dominance coefficient for all nonsynonymous mutations. However, such models are likely unrealistic, as *h* is thought to vary among different classes of deleterious mutations [14,15,17,18,25]. To systematically explore the parameter space for recessive deleterious mutations in humans, we examined the fit of models where *h* could differ for each bin of the discrete DFE. To focus on more biologically plausible models, we assumed that neutral mutations (|*s*|<1e-5) are additive and that all the other mutations could have *h* between 0 and 0.5. These constraints led to a total of 4096 *h-s* models to be tested. For each model, we considered values of *h* for each bin including 0.0, 0.05, 0.10, 0.15, 0.25, 0.35, 0.45, and 0.50. For a given combination of *h* values, we then inferred the proportions of mutations in each bin of the discrete DFE that maximizes the log-likelihood.

We found that these 4096 dominance models produced a wide range of DFE estimates, many exhibiting a very poor fit to the data (**Fig. 2**, top row). After removing a total of 3793 models that were significantly different than the MLE (1.92 LL units lower than the best model’s LL=- 1451.68), 303 models remained with good fit to the nonsynonymous SFS (hereafter, “high LL models”). These models demonstrate that a range of *h* values for each bin, ranging from additive to fully recessive, can yield a good fit to the nonsynonymous SFS (**Fig. 2**, middle row). However, the data constrains some of the parameter space. For example, models with *h*<0.15 for moderately deleterious mutations (10^!#^< |𝑠| ≤ 10^!$^) do not fit the data well, suggesting that such mutations are either additive or only partially recessive. Strongly deleterious mutations (10^!%^ < |𝑠| ≤ 0.5) can be more recessive, as models with *h*=0.05 fit the data (**Fig. 2**). Consistent with the data rejecting highly recessive models (*h*<0.05), we found that a highly recessive *h-s* relationship that was previously inferred for *Arabidopsis* by Huber et al. [25] also did not yield a good fit to human data (**Fig. S3**). This finding suggests that the parameters of the *h-s* relationship may greatly differ across species.

**Figure 2.**
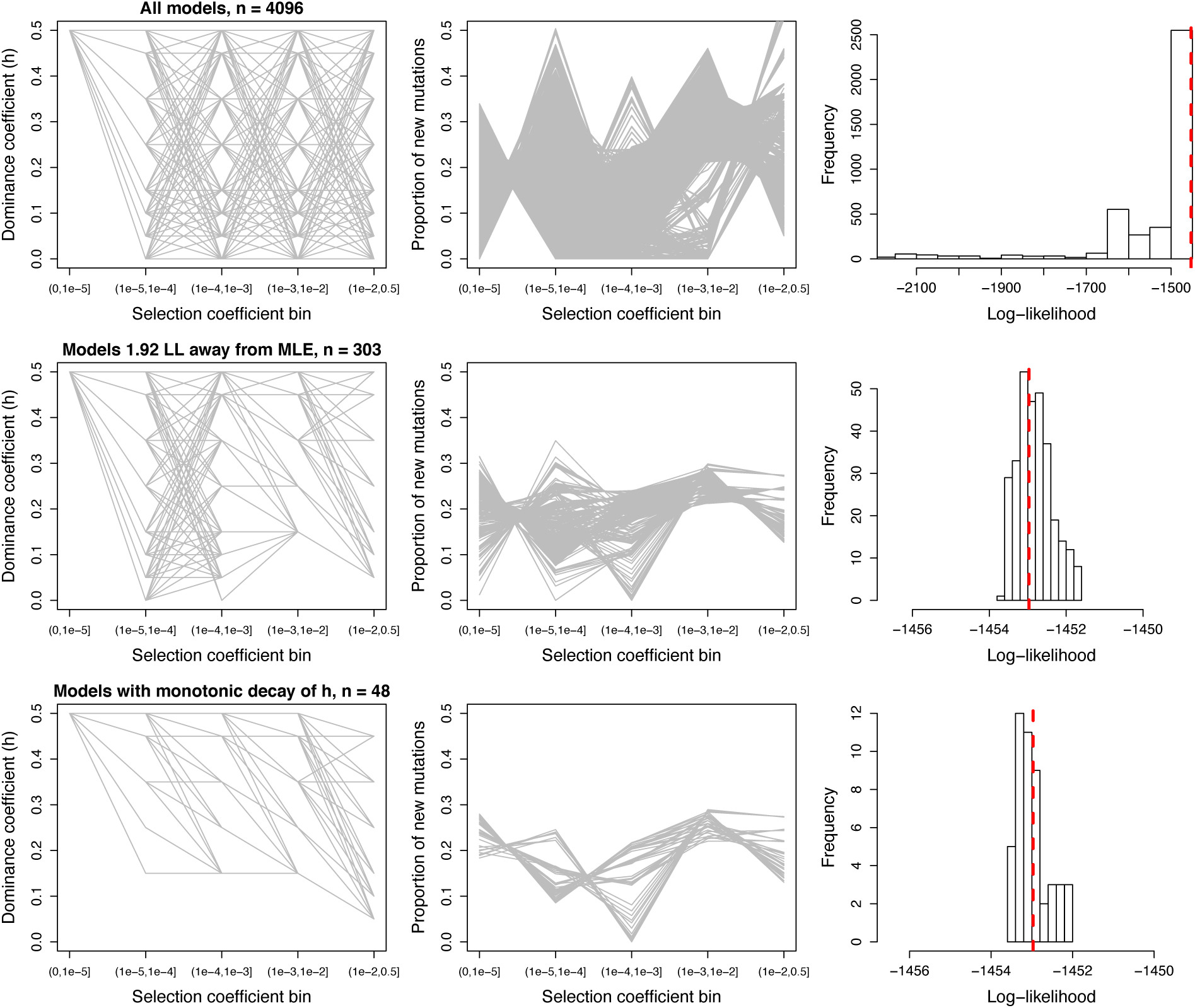
Visualizing DFE and dominance parameters for models with varying *h* for different discrete bins of *s*. Left column: Representation of *h-s* relationship for each model. Each model is shown as a gray line. Middle column: The resulting DFE inferred assuming each *h-s* relationship. Each model is shown as a gray line. Right column: The distribution of log-likelihood values when fitting different *h-s* discrete models. Red dashed line corresponds to the log-likelihood under an additive discrete model with 4 parameters (-1452.97). Top: All *h-s* models considered. Middle: Only those *h-s* models having a log-likelihood<1.92 units from that of the MLE are shown. Bottom: Only those *h-s* models having a log-likelihood<1.92 units from that of the MLE and that have a monotonic relationship between *h* and *s* are shown.

As another way of visualizing the impact of recessive mutations on model fit, we plotted the change in log-likelihood relative to a fully additive model while assuming progressively more recessive dominance coefficients for each selection coefficient bin of the DFE. In other words, we varied *h* for each bin of the DFE one-by-one while assuming *h*=0.5 for all other bins and determined the effects on model fit. We found that model fit changed minimally while varying *h* for the nearly neutral bin, whereas much greater changes in model fit were observed for other bins of *s* (**Fig. 3**). Specifically, models with *h*=0.0 for the weakly deleterious and moderately deleterious bins are ∼60 and ∼200 LL units worse than the fully additive model, respectively (**Fig. 3**). This suggests that, although some models with fully recessive weakly deleterious mutations can fit the data (**Fig. 2**), the constraint that all other bins are additive yields a poor fit (**Fig. 3**). Finally, for the strongly deleterious bin, we observe a loss of ∼13 LL units when *h*=0.0, though find that the fit is slightly improved when *h*=0.05 relative to the fully additive model (**Fig. 3**). This result is consistent with our above finding that strongly deleterious mutations appear to have a lower bound of *h* of 0.05 (**Fig. 2**).

**Figure 3:**
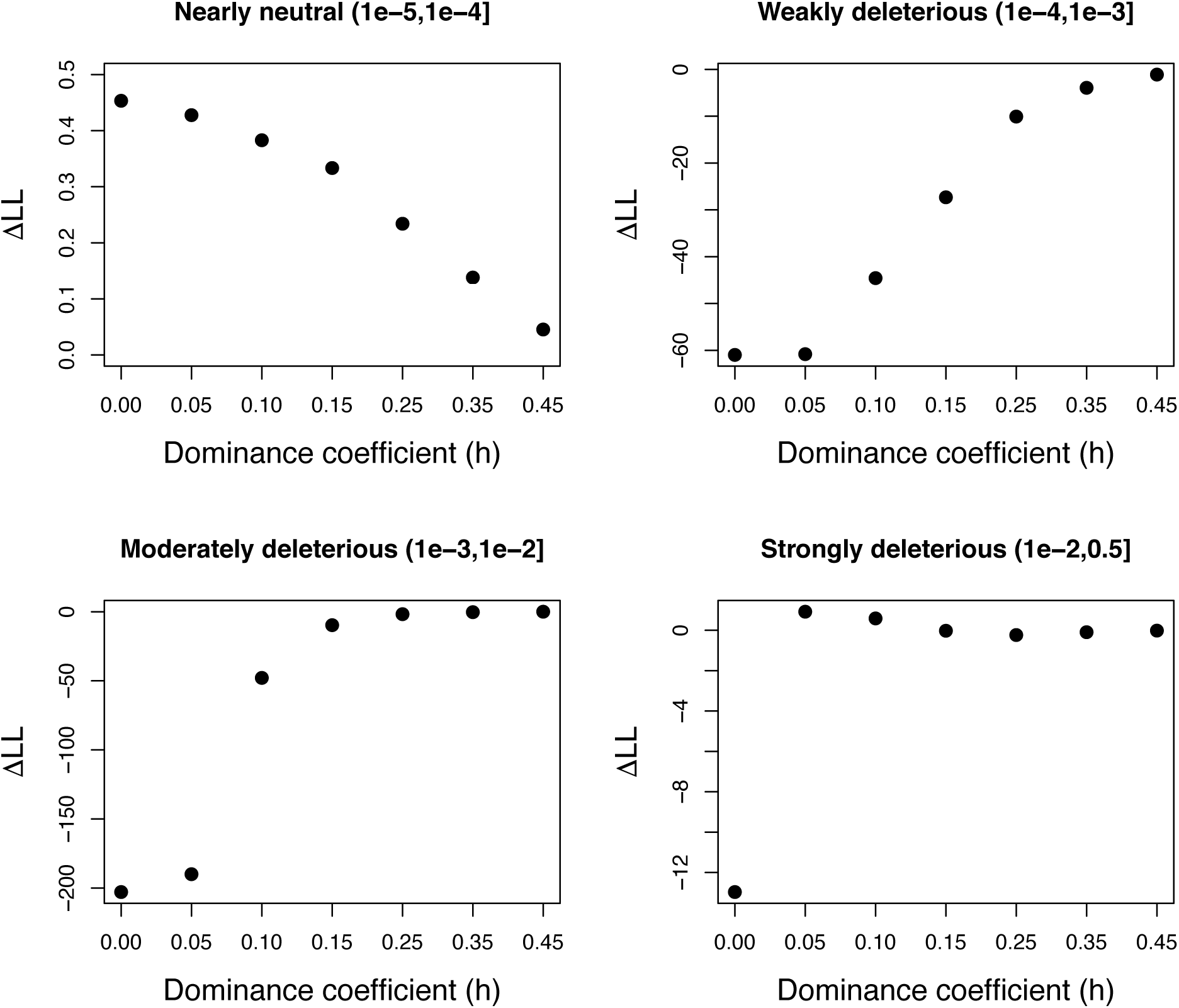
Exploring the impact of changing *h* for each selection coefficient bin under a discrete DFE model. Each plot shows the change in log-likelihood (ΔLL) relative to a model where all bins are assumed to be additive (*h*=0.5; LL=-1452.97). In each case, the dominance coefficient for the specified bin of the DFE (shown in each panel) was changed to a more recessive value (shown on the x-axis) while holding all other bins to *h*=0.5. Note that the model fit changes minimally as *h* becomes more recessive with the exception of making the weakly or moderately deleterious bins recessive. Strongly deleterious mutations show a complex pattern, where a model of *h*=0.05 results in a slight improvement in fit compared to the additive case while a fully recessive model (*h*=0) fits worse.

### Testing models with an *h-s* relationship

Because experimental and molecular population genetics work has previously suggested that more deleterious mutations tend to be more recessive [14,17,18,25], we next restricted all the high LL models (n=303) to those with a monotonic decay in their dominance coefficients from the neutral to the strongly deleterious mutation classes. A total of 48 high LL models with a monotonic decay remained, with log-likelihoods ranging from -1452.02 to -1453.50 (**Fig. 2**, bottom row). Under these conditions, nearly neutral, weakly deleterious, and moderately deleterious mutations have a lower bound dominance coefficient of *h*=0.15, while strongly deleterious mutations have a lower bound *h*=0.05. Across these different models, the inferred discrete DFE proportions remain relatively consistent for some selection coefficient bins, though are variable for others. For instance, the inferred proportions for the moderately deleterious bin remained within 0.22-0.29, whereas the strongly deleterious bin varied more widely between 0.13-0.27 (**Fig. 2**).

To obtain estimates of *h* and the DFE while accounting for model uncertainty, we next conducted model averaging using the 𝛥AICs as weights of contributions to the parameter estimation (see **Methods**). When averaging across all 4096 models, we estimate an overall *h* of 0.29, including an estimate of *h*=0.23 for strongly deleterious mutations (**Table 1**). Model averages when using high LL models (n=303) and monotonic high LL models (n=48) both suggest an overall *h* of ∼0.35 along with DFEs that are highly similar (**Table 1**). However, the estimated *h* values for each bin of the DFE vary substantially, with an *h* of 0.20 for strongly deleterious mutations when using the high LL models compared to *h*=0.13 for strongly deleterious mutations when only including monotonic models (**Table 1**).

**Table 1:**
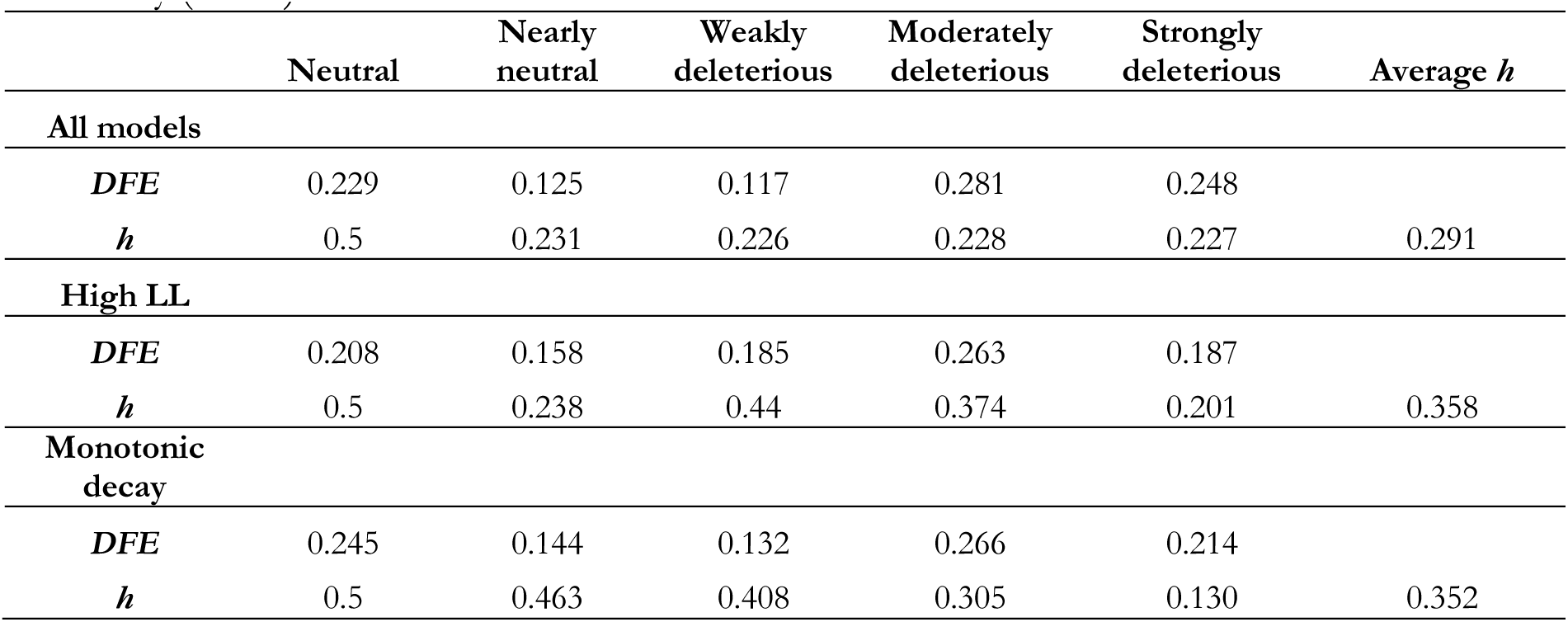
Summary of model averaging results. . Results are shown when considering all possible models (n=4096), high LL models within 1.92 LL units of the MLE (n=303), and high LL models with a monotonic decay in *h* (n=48). For each model, parameters of the discrete DFE (i.e. the proportions of mutations falling in each bin) and corresponding *h* for each bin of the DFE are shown. Selection coefficient bins are defined as: neutral (0 < |𝑠| ≤ 10^!"^), nearly neutral (10^!"^ < |𝑠| ≤ 10^!#^), weakly deleterious (10^!#^ < |𝑠| ≤ 10^!$^) moderately deleterious (10^!$^ < |𝑠| ≤ 10^!%^), and strongly deleterious (10^!%^ < |𝑠| ≤ 0.5). Note that models were constrained to enforce additivity (*h*=0.5) for neutral mutations.

| | Neutral | Nearly neutral | Weakly deleterious | Moderately deleterious | Strongly deleterious | Average $h$ |
| --- | --- | --- | --- | --- | --- | --- |
| **All models** |  |  |  |  |  |  |
| *DFE* | 0.229 | 0.125 | 0.117 | 0.281 | 0.248 |  |
| *h* | 0.5 | 0.231 | 0.226 | 0.228 | 0.227 | 0.291 |
| **High LL** |  |  |  |  |  |  |
| *DFE* | 0.208 | 0.158 | 0.185 | 0.263 | 0.187 |  |
| *h* | 0.5 | 0.238 | 0.44 | 0.374 | 0.201 | 0.358 |
| **Monotonic decay** |  |  |  |  |  |  |
| *DFE* | 0.245 | 0.144 | 0.132 | 0.266 | 0.214 |  |
| *h* | 0.5 | 0.463 | 0.408 | 0.305 | 0.130 | 0.352 |

### Simulating deleterious variation in human populations under a range of dominance models

Our results suggest a range of dominance models can fit human genetic variation data, though with support for an overall mean *h* for nonsynonymous mutations of ∼0.35 and evidence for the possibility of an *h-s* relationship. In light of these findings, our next aim was to explore how these estimates of dominance parameters may influence models of deleterious variation and genetic load across human populations. Specifically, we sought to revisit the question of how the out-of-Africa bottleneck may have impacted the relative burden of deleterious variation in African and non-African human populations. As previous studies have demonstrated that dominance is a key component influencing potential differences in deleterious burden [5–9], reevaluating these patterns with dominance models that are fit to human genetic variation data may provide further clarity on this topic.

To explore the relative burden of deleterious variation in African and non-African human populations behaves under our estimated models of dominance, we ran forward-in-time simulations of deleterious genetic variation using SLiM [33–35] under a human demographic model [36] employing a range of dominance models as suggested by our results. These models include a “Weakly Recessive” model with average *h*=0.40 and *h*=0.15 for strongly deleterious mutations, a “Moderately Recessive” model with average *h*=0.34 and *h*=0.10 for strongly deleterious mutations, and a “Strongly Recessive” model with average *h*=0.25 and *h*=0.05 for strongly deleterious mutations (**Table 2**). We chose these models to encompass the range of inferred dominance parameters from our results while also exploring plausibly low dominance coefficients for strongly deleterious mutations, given the widespread evidence for strongly deleterious mutations being highly recessive [14,15,17,25,37,38]. Note that the above models were selected from the broader set of 48 high LL models with a monotonic decay (**Fig. 2**; **Table 2**); thus, these models all exhibit similarly good fit to the human nonsynonymous SFS.

**Table 2:** Summary of DFE & dominance models used for simulations. For each model, parameters of the discrete DFE and corresponding *h* for each bin of mutations are shown. Selection coefficient bins are defined as: neutral (0 < |𝑠| ≤ 2 ∗ 10^!"^), nearly neutral (2 ∗ 10^!"^ < |𝑠| ≤ 2 ∗ 10^!#^), weakly deleterious (2 ∗ 10^!#^ < |𝑠| ≤ 2 ∗ 10^!$^) moderately deleterious (2 ∗ 10^!$^ < |𝑠| ≤ 2 ∗ 10^!%^), and strongly deleterious (2 ∗ 10^!%^ < |𝑠| ≤ 1).

| | Neutral | Nearly neutral | Weakly deleterious | Moderately deleterious | Strongly deleterious | Average $h$ |
| --- | --- | --- | --- | --- | --- | --- |
| **Strongly recessive** |  |  |  |  |  |  |
| **DFE** | 0.201 | 0.222 | 0.018 | 0.286 | 0.274 |  |
| **$h$** | 0.5 | 0.45 | 0.25 | 0.15 | 0.05 | 0.26 |
| **Moderately recessive** |  |  |  |  |  |  |
| **DFE** | 0.244 | 0.152 | 0.125 | 0.259 | 0.22 |  |
| **$h$** | 0.5 | 0.5 | 0.45 | 0.25 | 0.1 | 0.34 |
| **Weakly recessive** |  |  |  |  |  |  |
| **DFE** | 0.259 | 0.125 | 0.176 | 0.254 | 0.186 |  |
| **$h$** | 0.5 | 0.5 | 0.5 | 0.35 | 0.15 | 0.40 |

For these simulations, we modelled coding regions for 22 autosomes, yielding results for ∼30Mb of total sequence (see **Methods**). Under each dominance model, we outputted the predicted genetic load at the conclusion of the simulation, which measures the reduction in mean population fitness due to segregating and fixed deleterious mutations [4,39]. Additionally, we also outputted the predicted inbreeding load under each model, which measures the potential severity of inbreeding depression (i.e., the quantity of recessive deleterious variation that is concealed as heterozygotes) in a population [3,39,40]. The inbreeding load (often referred to as the ‘number of lethal equivalents’ or *B*) therefore provides a complementary perspective on the burden of deleterious variation.

Moreover, several empirical inbreeding load estimates exist for humans, suggesting a range for *B* between ∼0.7-2.5 [40,41]. Thus, comparing these empirical estimates to those predicted by each dominance model can serve as an additional source of evidence to validate dominance and selection parameters.

Our simulation results suggest that the predicted patterns of load depend somewhat on the dominance model employed. Specifically, in the Weakly and Moderately Recessive models, we observe a slight increase in genetic load in the non-African population (1.7% and 1.0% increase, respectively), whereas in the Strongly Recessive model, we observe a 1.2% decrease in genetic load in non-African populations (**Fig. 4A**). By contrast, although the Weakly Recessive model suggests a 2% increase in the inbreeding load in non-African populations, the Moderately and Strongly Recessive models estimate a slight ‘purging’ of the inbreeding load due to the out-of-Africa bottleneck (1.7% and 1.8% decrease, respectively; **Fig. 4B**). Moreover, the total inbreeding load predicted varies by model, with B=∼0.55 predicted for the Weakly Recessive model, B=∼0.92 predicted for the Moderately Recessive model, and B=∼2.1 predicted for the Strongly Recessive model (**Fig. 4B**). Notably, this range of predicted inbreeding loads is in good agreement with the range of empirical estimates in humans of ∼0.7-2.5 [40,41]. Finally, the counts of derived nonsynonymous alleles showed a general trend of a small increase in non-African populations, with 1% increase under the Weakly Recessive model, a 1.8% increase under the Moderately Recessive model, and a 0.5% increase under the Strongly Recessive model (**Fig. 4C**). Notably, these trends were not entirely predictive of the patterns observed for the genetic load (**Fig. 4A**).

**Figure 4:**
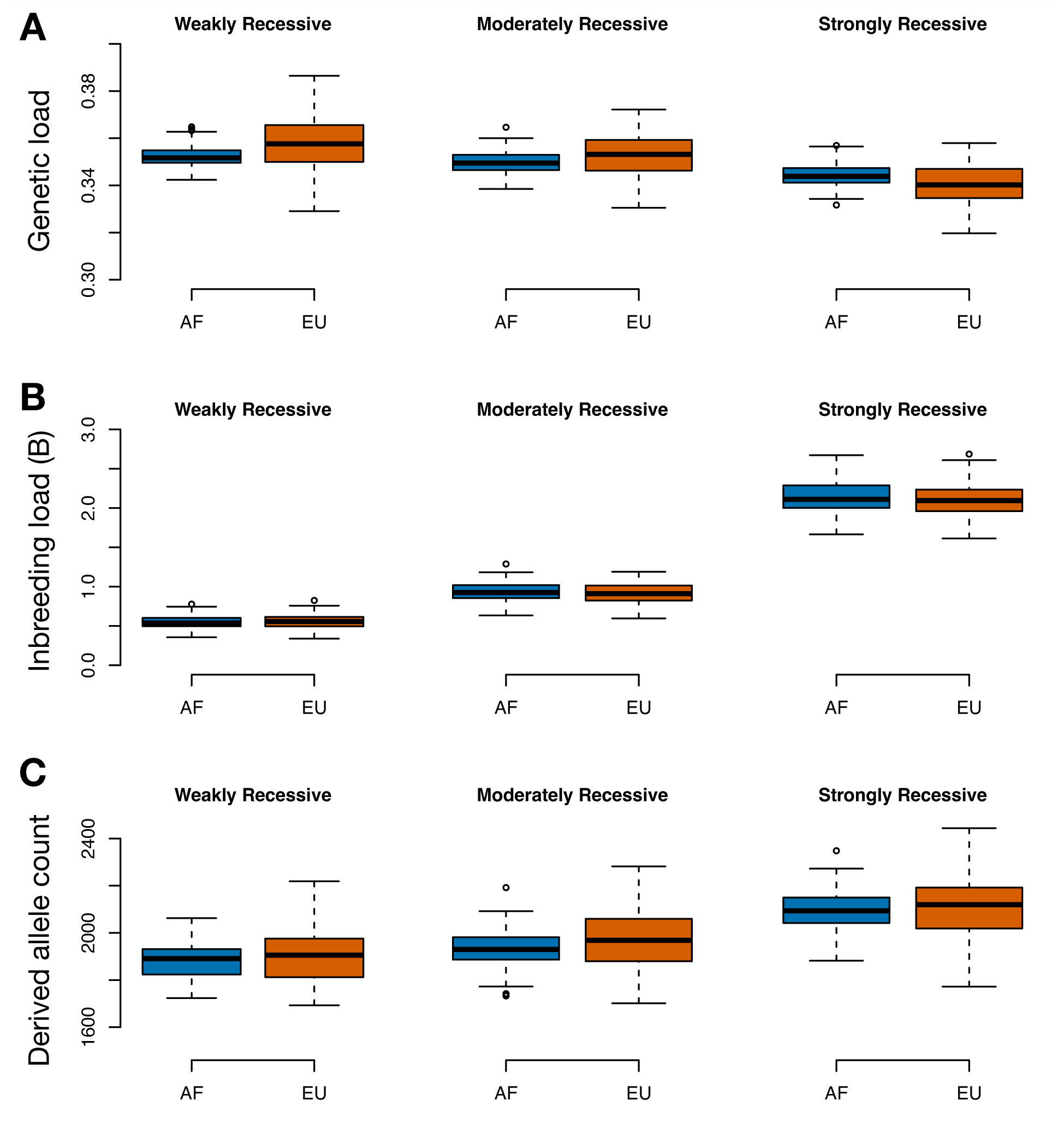
Simulation results comparing predicted genetic load, inbreeding load, and derived allele count for African (AF) and European (EU) populations under four different DFE and dominance models. (A) Predicted genetic load in African and European populations. (B) Predicted inbreeding load in African and European populations. Note that empirical estimate of *B* in humans range from ∼0.7-2.5 [40,41]. (C) Predicted derived deleterious allele count in African and European populations. Results are shown as boxplots summarizing output from 100 simulation replicates under each DFE and dominance model. See Table 2 for details on each DFE and dominance model.

## Discussion

Here, we have investigated the fit of dominance and selection models for nonsynonymous mutations humans and explored implications for the relative burden of deleterious variation in African and non-African human populations. Our results demonstrate that a wide range of dominance models are consistent with patterns of nonsynonymous genetic variation in humans, including models with a strong *h-s* relationship. Although our analysis is unable to fully overcome issues of identifiability for *h* and *s* parameters, some general conclusions can be drawn. First, we demonstrate that highly recessive models with a single dominance coefficient (*h*<0.15) cannot yield a good fit to the nonsynonymous SFS in humans (**Fig. 1; Figs. S1-S2**). Next, we find that many models with an *h-s* relationship yield a good fit to the data (**Fig. 2**), as predicted by previous theoretical and empirical work [14,17,19,25,42]. Additionally, through model averaging, we estimate an average *h* on the order of ∼0.35, though with much greater uncertainty for *h* values of individual selection coefficient bins (**Fig. 2**; **Table 1**). Notably, we find that the *Arabidopsis h-s* relationship parameters estimated by Huber et al. [25] do not fit human data (**Fig. S3**), suggesting that dominance parameters are likely to vary across species. Finally, we also find that that models with *h*=0.05 for strongly deleterious mutations provide a small improvement in fit to the SFS (**Fig. 3**), consistent with previous work demonstrating that such mutations are likely to be highly recessive [14,15,17,25,37,38]. For instance, the impacts of recessive lethal mutations in humans are well documented [37,43,44], though evidence from *Drosophila* suggests that such lethal mutations may in fact have a small fitness consequence in the heterozygote state [17], consistent with our findings (**Figs. 2 & 3**) Dominance is an essential determinant of the influence of demography on patterns of deleterious variation [1,2,4,5]. In humans, previous work has found that dominance plays a key role in determining the impact of the out-of-Africa bottleneck on relative patterns of deleterious variation and genetic load in African and non-African populations [5–9]. In the absence of estimates of dominance parameters in humans, previous studies have typically assumed the extreme cases of additive and fully recessive models [8–10] or used *ad hoc* dominance parameters [6]. Our simulation results under a range of plausible dominance models fit to data further demonstrate the subtle influence of dominance on the relative burden of genetic load in African and non-African populations. Specifically, we find that genetic load may be slightly elevated in non-African populations when deleterious mutations are weakly or moderately recessive, whereas genetic load in non-African populations may in fact be slightly diminished if deleterious mutations are more strongly recessive (**Fig. 4**). This diminished genetic load in non-African populations in the Strongly Recessive model is driven by a slight purging of the inbreeding load during the out-of-Africa bottleneck (**Fig. 4**), a process that is known to be most efficient for highly recessive strongly deleterious mutations [1–3]. Finally, these simulation results also help to further validate our dominance models, demonstrating that selection and dominance models estimated from genomic variation datasets predict inbreeding loads that are broadly consistent with empirical measures.

Specifically, these models predict a *B* between ∼0.5-2.1, a range that is strikingly similar to that suggested by empirical studies of ∼0.7-2.5 [40,41]. This result helps bridge the gap between molecular studies of dominance and selection parameters and more direct measures of fitness in humans, suggesting that results from these very different approaches can be reconciled. Moreover, this result also suggests that nonsynonymous mutations alone can account for much of the inbreeding depression observed in humans.

Our work also has implications for studies of deleterious variation in non-human taxa. In particular, there has been a great deal of recent interest in modelling the impact of recessive deleterious variation on extinction risk in small and isolated populations [2,39,45–47]. Our analysis can help guide such studies by informing the parameterization of dominance and selection models, as many of these studies are focused on endangered species of mammals that may have similar population genetic parameters to humans [39]. For instance, it has been previously suggested that the historical population size of a species may greatly influence risk of extinction due to inbreeding depression, particularly in cases where strongly deleterious alleles (1e-2<|*s*|=<1) are highly recessive (*h*<0.05) [1–3]. The results of our simulation analysis indicate that models with fully recessive strongly deleterious mutations are not compatible with empirical estimates of the inbreeding load in humans (**Fig. 4**), suggesting that this relationship between demography and deleterious variation may be somewhat dampened, at least in humans. Instead, we find that an *h* for strongly deleterious mutations on the order of ∼0.05-0.15 could better explain empirical estimates of the inbreeding load (**Fig. 4**). However, we note that a major limitation of our study is that we are unable to finely estimate selection and dominance parameters for strongly deleterious mutations, which as defined here encompass a wide range of |*s*| from 0.01 to 1. This limitation is due to SFS-based methods being underpowered for estimating the strongly deleterious tail of the DFE [44], due to the fact that such mutations tend not to be segregating in genetic variation datasets [48–50]. Thus, an important area for future work is to further refine selection and dominance parameters for strongly deleterious mutations and determine whether dominance parameters may differ between strongly deleterious mutations (|*s*| on the order of ∼0.01) and lethal mutations (|*s*|=1).

Our study also remains limited by an inability to statistically favor one dominance model over another. For example, an additive model continues to fit the SFS very well, exhibiting similar fit to more complex recessive models (**Figs. 1-2; Tables S1-S2**). To overcome this, we use a model averaging approach (**Table 1**), which highlights some general trends, though does not necessarily indicate that complex models with recessive mutations are an improvement over an additive model. Nevertheless, the plausibility of these more complex models with highly recessive strongly deleterious mutations are supported by previous theoretical and experimental work on dominance in non-human taxa [14,17,19,25,42] as well as the broad literature on recessive disease and inbreeding depression in humans [37,40,41,51–56]. Although we remain unable to determine an optimal dominance model in humans, our work provides a range of useful models that can be employed in future analyses [57]. To improve on our findings and obtain more precise estimates of dominance in humans, some recent work has suggested that there may be information on dominance from patterns of linkage disequilibrium [58,59] or from patterns of transmission in pedigrees [60].

Alternatively, leveraging genomic datasets from populations with varying levels of inbreeding in humans or other mammals, as done by Huber et al. [25] in *Arabidopsis*, may also offer a fruitful avenue for research. Future studies should continue to explore these and other avenues for inferring dominance parameters to better inform our understanding of the evolutionary significance of dominance.

## Materials and Methods

### Data

We downloaded SNP genotype data for 432 individuals with European ancestry from the 1000 Genomes Project phase 3 release [31] from http://ftp.1000genomes.ebi.ac.uk/vol1/ftp/release/20130502/. We followed filtering steps as described in Kim et al. [27]. Specifically, only unrelated individuals were used and only sites from exome-targeted sequencing that passed the strict mask filter criteria (as defined in [31]) were kept. We then extracted the synonymous and nonsynonymous sites based on the 1000 Genomes Project- filtered annotations. The folded site frequency spectra were computed by tabulating the observed counts of the minor allele frequencies for synonymous and nonsynonymous variants separately. The synonymous and nonsynonymous site frequency spectra were then used for the inference of demography and DFE, respectively. We also computed the length of synonymous (LS) and the length of nonsynonymous sites (LNS) sites which were used for the estimation of population genetic parameters (see below; **Table S2**).

### Demographic Parameters

Because demographic history distorts allele frequencies in similar ways to selection, we used the synonymous SFS to tease apart the effect of demography and selection on the SFS. Specifically, we inferred the parameters in a three-epoch demographic model (out-of-Africa European demographic model with the occurrence of a bottleneck, a recovery period, followed by a recent exponential population growth) from the 1000 Genomes Project European synonymous SFS using ∂a∂i [61]. We computed changes in population size relative to the ancestral population size (*Nanc*) by storing the maximum-likelihood estimates (MLEs) of demography and population sizes after 30 iterations (MLEs are shown in **Table S1**). The mutation rate at synonymous sites (*θ*S) was then estimated as the scaling factor difference between the optimized SFS and the empirical data using the function *dadi.Inference.optimal_sfs_scaling* with ∂a∂i [61]. The ancestral population size *Nanc* was obtained from the inferred scaled mutation rate at synonymous sites (*θ*S) using the formula: 𝜃_(_ = 4𝑁_)*+_ ∗ 𝜇 ∗ 𝐿_(_, where 𝐿_!_ is the is total count of synonymous sites, and 𝜇 is the per-base-pair mutation rate. We assumed a mutation rate of 1.5e-8 mutation per site per generation [62]. To compute the nonsynonymous scaled mutation rate (*θ*NS) we assumed a 2.31 ratio between 𝐿_"!_ and 𝐿_!_ [32]. All computed population genetic parameters are described in **Table S2**.

### DFE inference under a gamma model with varying *h*

To infer selection while accounting for demography, we assumed the estimated demographic parameters obtained from the synonymous SFS (**Table S1**), and used the nonsynonymous SFS to infer the DFE with Fit∂a∂i using a Poisson likelihood function as described in [26,27,32]. In Fit∂a∂i, fitness is parameterized such that the mutant homozygote has fitness 1-2*s* and the heterozygote has fitness 1-2*sh*. Thus, selection coefficients for deleterious mutations are positive and range from 0 to 0.5, where 0.5 is considered a lethal mutation with fitness of 0.

Our initial aim was to fit a gamma distributed DFE to the nonsynonymous SFS under varying *h* values including 0.0, 0.05, 0.10, 0.15, 0.25, 0.35, 0.45, 0.50, 0.75, and 1.0. The gamma distribution has two parameters (shape and scale, often denoted 𝛼 and 𝛽) and has previously been shown to provide a reasonable distribution for the DFE [26–29]. To determine the expected SFS for nonsynonymous mutations under varying values of *h*, we used the equation in Williamson et al. [23] implemented within ∂a∂i in the function *phi_1D* to compute the quasi-stationary distribution of allele frequency for a given site under the assumptions of the Wright-Fisher model (random mating, constant population size, non-overlapping generations):

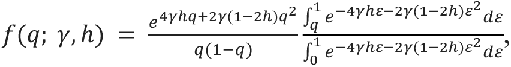

where *q* is the frequency of the derived nucleotide in the population, 𝛾 is the scaled selection coefficient (*s*\*2*Nanc*), and *h* is the dominance coefficient. We then used the *Integration.one_pop* function to update 𝑓(𝑞; 𝛾, ℎ) to become the transient distribution of allele frequencies after the population changed size. By assuming that sites are independent, we can expand the above formula to multiple sites by simply assuming that the allele frequency of a given mutant allele is a random draw from the above distribution.

Each entry of the SFS, 𝑥_4_, is the count of the number of variants at which the derived nucleotide present 𝒊 times in a sample size of 𝑛 individuals, for 𝒊 1, 2, . . ., 𝑛 − 1. Then the expected value for each entry of the SFS vector is 𝜃𝐹(𝑛, 𝑖; 𝛾, ℎ) [23]:

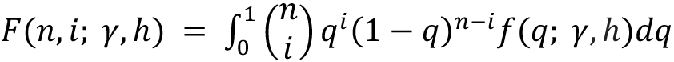

where 𝜃 is the per generation mutation rate of the sampled region. If mutations enter a population in each generation following a Poisson distribution [22], then each entry of the SFS (𝑥_4_) is expected to be Poisson distributed [22]. Given the full probability distribution of each entry of the SFS, the model parameters can be estimated in a maximum-likelihood framework within Fit∂a∂i.

We used this approach as implemented in Fit∂a∂i to compute the expected nonsynonymous SFS over a grid of 1000 log-spaced values of the population-scaled selection coefficient 𝛾 (*s*\*2*Nanc*). To maintain computational feasibility and avoid issues of numerical instability with the diffusion approximation due to large *s* [63], we restricted the range of *s* to 1e-5 to 0.25, thus assuming that mutations with |s|>0.25 were not segregating in the 1000G dataset. We then inferred the shape and scale parameters of the gamma distribution from 25 runs of Fit∂a∂i by integrating over the grid of expected SFS and fitting model output to the empirical nonsynonymous SFS. Parameters with the highest Poisson log-likelihood were chosen. For plotting, the scale parameter was divided by 2*Nanc* to no longer be scaled by the ancestral population size. Details of parameters used for inference and rescaling are provided in **Table S2.**

### Inference under a discrete DFE model with varying *h*

In addition to testing a gamma distribution for the DFE, we also examined the fit of the discrete DFE under varying dominance coefficients. The discrete DFE can be seen as a mixture of uniform distributions consisting of five bins defined based on a range of selection coefficients. The bins were defined as: neutral (0 < |𝑠| ≤ 10^!"^), nearly neutral (10^!"^ < |𝑠| ≤ 10^!#^), weakly deleterious (10^!#^ < |𝑠| ≤ 10^!$^) moderately deleterious (10^!$^< |𝑠| ≤ 10^!%^), and strongly deleterious (10^!%^ < |𝑠| ≤ 0.5). The proportions of new mutations in each bin were the parameters inferred using Fit∂a∂i. Note that, even though there are 5 bins of the DFE, the constraint that all 5 bins must sum to 1 means that only 4 parameters are actually inferred from the data. To estimate these parameters, we used the same approach as outlined above for the gamma distribution.

### Inference under a discrete DFE with multiple dominance coefficients

To better explore the fit of models with recessive mutations, we tested models where each bin of the discrete DFE could have its own value of *h* including 0.0, 0.05, 0.10, 0.15, 0.25, 0.35, 0.45, 0.50. To constrain the set of possible models, we assumed that neutral mutations (0 < |𝑠| ≤ 10^!"^) were additive (*h*=0.5) and tested out all combinations of *h* for the four other bins of the discrete DFE. This resulted in a total of 4096 (=8^4^, where 8 is the number of *h* values tested and 4 is the number of bins of the DFE) models being fit to the nonsynonymous SFS.

To fit these models, we used the same approach outlined above including estimating a demographic model using the synonymous SFS and then estimating the parameters of the discrete DFE using the nonsynonymous SFS. For each bin of the discrete DFE, we used the precomputed expected SFS under the assumed value of *h* for the given range of *s*. For instance, in the case of a model with *h*=0.5 for neutral mutations, *h*=0.35 for nearly neutral mutations, *h*=0.25 for weakly deleterious mutations, *h*=0.15 for moderately deleterious mutations, and *h*=0.05 for strongly deleterious mutations, we created a grid of expected nonsynonymous SFS using expectations for *h*=0.5 for the *s* range from 0 to 1e-5, *h*=0.35 for the *s* range from 1e-5 to 1e-4, *h*=0.25 for the *s* range from 1e-4 to 1e-3, *h*=0.15 for the *s* range from 1e-3 to 1e-2, and *h*=0.05 for the *s* range from 1e-2 to 0.5. To maintain computational feasibility, we ran only 5 Fit∂a∂i runs for each of the 4096 dominance models and picked the parameters from the run with the highest Poisson log-likelihood.

### Computing AIC

To compare non-nested DFE models, such as the gamma and discrete models, and estimate parameters of interest using a model averaging approach (see below), we first transformed the log likelihood using the Akaike information criteria:

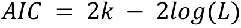

where 𝑘 is the number of estimated parameters (2 for the gamma DFE and 4 for the discrete DFE models) and 𝐿 is the maximum likelihood of each model estimated. The preferred AIC model is the one with the minimum AIC value. These AIC values also were then used as weights of the model averaging (below).

### Model averaging with AIC

Because multiple *h-s* relationship models fit the data, we also used a model averaging approach to estimate the DFE and dominance parameters [64,65]. Let *x* refer to the parameter of interest (e.g. all entries in the discretized DFE and their respective dominance coefficients), 𝑥_4_ refer to the MLE under model 𝑖 that has an AIC value of 𝐴𝐼𝐶_4_. In this approach, the parameter of interest,

𝑥 is estimated. Then, the model average estimate (𝑥_)67_) accounting for the contribution of each model is averaged using Akaike weights:

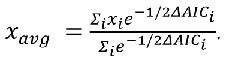

The 𝛥𝐴𝐼𝐶_4_ is obtained as:

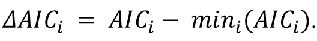

We computed model averages first considering all possible discrete models with varying *h* (**Fig. 2**, top row), next considering only models that were 1.92 LL units away from the MLE (**Fig. 2**, middle row), and finally considering only models that were 1.92 LL units away from the MLE and had a monotonic decay in *h* (**Fig. 2**, bottom row).

### Simulation methods

We ran forward-in-time simulations under the Wright-Fisher model using SLiM v4.0.1 [33–35] employing DFE and dominance parameters inferred above to revisit the question of how the out-of-Africa bottleneck has influenced the relative burden of deleterious variation in African and non-African human populations. We modelled deleterious alleles occurring as nonsynonymous mutations in coding sequence, with 22,500 genes on 22 autosomes each 1340 bp in length, resulting in a total sequence length of 30.16 Mb [66]. Following Robinson et al. [12], we assumed a recombination rate of 1e-3 crossovers per site per generation between genes, 0.5 between chromosomes, and no recombination within genes. The mutation rate for deleterious alleles was set to 1.5e-8*2.31/3.31=1.05e-8 mutations per site per generation, where 1.5e-8 is the genome-wide mutation rate in humans [62] and 2.31/3.31 is the fraction of coding mutations that are assumed to be nonsynonymous [32]. We set the demographic parameters using the two-population model from Tennessen et al. [36], inferred from large samples of individuals of African and European descent. Specifically, this model assumes an ancestral population size of *Ne*=7,310, followed by growth in Africa to *Ne*=14,474, with the European population diverging 2,040 generations before present and experiencing a bottleneck at *Ne*=1,861 for 1,120 generations, followed by exponential growth over the last 204 generations in both populations to *Ne*=424,000 in Africa and *Ne*=512,000 in Europe.

We employed three different DFE and dominance models in our simulations (**Table 2**), chosen to encompass the range of parameters observed in our monotonic *h-s* relationship results while also modelling strongly deleterious mutations as being highly recessive (*h*<=0.15) as informed by studies in other taxa [14,15,17,25]. These models include a “Weakly Recessive” model with average *h*=0.40 and *h*=0.15 for strongly deleterious mutations, a “Moderately Recessive” model with average *h*=0.34 and *h*=0.10 for strongly deleterious mutations, and a “Strongly Recessive” model with average *h*=0.26 and *h*=0.06 for strongly deleterious mutations (**Table 2**). Note that these models were selected from the broader set of 48 high LL models with a monotonic decay (**Fig. 2**; **Table 1**) and therefore all exhibit a similar fit to the human nonsynonymous SFS. Finally, also note that, unlike Fit∂a∂i, fitness in SLiM parameterized such that the mutant homozygote has fitness 1-*s* and the heterozygote has fitness 1-*sh*. Thus, the selection coefficient bins for these SLiM simulations were defined as: neutral (0 ≤ |𝑠| < 2 ∗ 10^!"^), nearly neutral (2 ∗ 10^!"^ ≤ | 𝑠| < 2 ∗ 10^!#^), weakly deleterious (2 ∗ 10^!#^ ≤ |𝑠| < 2 ∗ 10^!$^) moderately deleterious (2 ∗ 10^!$^ ≤ |𝑠| < 2 ∗ 10^!%^), and strongly deleterious (2 ∗ 10^!%^ ≤ | 𝑠| ≤ 1).

For each DFE and dominance model, we ran 100 simulation replicates and, at the conclusion of the simulation, outputted the mean genetic load, mean inbreeding load, and average number of derived deleterious alleles per individual in each population, taken from a sample of 100 individuals. Here, we define genetic load as the reduction in fitness due to segregating and fixed deleterious alleles, where fitness is multiplicative across sites [4,39,67]. We measure the inbreeding load as the “number of haploid lethal equivalents”, which quantifies the summed selective effects of heterozygous recessive deleterious alleles in a population [3,39,40]. Due to the high computational load of these simulations, we ran simulations for only two autosomes and projected results to a full genome of 22 autosomes. To do this, we exponentiated the outputted relative fitness values by 11 (subtracting this from 1 to obtain genetic load) and multiplied the outputted inbreeding load and derived allele counts by 11 (as these quantities are summed across chromosomes). Note that this procedure results in the simulation variance being overestimated, though the averages of these quantities are unbiased.

## Acknowledgements

We are grateful to Maria Izabel Cavassim Alves for her instrumental role in the early stages of this project and to Bernard Kim for assistance with Fit∂a∂i. C.C.K. and K.E.L. were supported by the National Institutes of Health (R35GM119856 to K.E.L.).

## Data Availability

All scripts are available on GitHub (https://github.com/ckyriazis/dominance).

## Author Contributions

Conceptualization: Christopher C. Kyriazis, Kirk E. Lohmueller. Formal analysis: Christopher C. Kyriazis, Kirk E. Lohmueller.

Funding acquisition: Kirk E. Lohmueller.

Investigation: Christopher C. Kyriazis, Kirk E. Lohmueller. Methodology: Christopher C. Kyriazis, Kirk E. Lohmueller. Project administration: Kirk E. Lohmueller.

Resources: Kirk E. Lohmueller.

Software: Christopher C. Kyriazis, Kirk E. Lohmueller. Supervision: Kirk E. Lohmueller.

Visualization: Christopher C. Kyriazis, Kirk E. Lohmueller.

Writing – original draft: Christopher C. Kyriazis, Kirk E. Lohmueller. Writing – review & editing: Christopher C. Kyriazis, Kirk E. Lohmueller.

## References

1. Hedrick PW. Lethals in finite populations. Evolution (N Y). 2002;56: 654–657.

2. Kyriazis CC, Wayne RK, Lohmueller KE. Strongly deleterious mutations are a primary determinant of extinction risk due to inbreeding depression. Evol Lett. 2021;5: 33–47. doi:10.1002/evl3.209

3. Hedrick PW, Garcia-Dorado A. Understanding inbreeding depression, purging, and genetic rescue. Trends Ecol Evol. 2016;31: 940–952. doi:10.1016/j.tree.2016.09.005

4. Kirkpatrick M, Jarne P. The effects of a bottleneck on inbreeding depression and the genetic load. Am Nat. 2000;155: 154–167. doi:10.1086/303312

5. Balick DJ, Do R, Cassa CA, Reich D, Sunyaev SR. Dominance of deleterious alleles controls the response to a population bottleneck. PLoS Genet. 2015;11: 1–23. doi:10.1371/journal.pgen.1005436

6. Henn BM, Botigué LR, Peischl S, Dupanloup I, Lipatov M, Maples BK, et al. Distance from sub-Saharan Africa predicts mutational load in diverse human genomes. Proc Natl Acad Sci. 2016;113: E440–E449. doi:10.1073/pnas.1510805112

7. Simons YB, Sella G. The impact of recent population history on the deleterious mutation load in humans and close evolutionary relatives. Curr Opin Genet Dev. 2016;41: 150–158. doi:10.1016/j.gde.2016.09.006

8. Do R, Balick D, Li H, Adzhubei I, Sunyaev S, Reich D. No evidence that selection has been less effective at removing deleterious mutations in Europeans than in Africans. Nat Genet. 2015;47: 126–131. doi:10.1038/ng.3186

9. Simons YB, Turchin MC, Pritchard JK, Sella G. The deleterious mutation load is insensitive to recent population history. Nat Genet. 2014;46: 220–224. doi:10.1038/ng.2896

10. Lohmueller KE, Indap AR, Schmidt S, Boyko AR, Hernandez RD, Hubisz MJ, et al. Proportionally more deleterious genetic variation in European than in African populations. Nature. 2008;451: 994–997. doi:10.1038/nature06611

11. Kim BY, Huber CD, Lohmueller KE. Deleterious variation shapes the genomic landscape of introgression. PLoS Genet. 2018;14: 1–30.

12. Robinson JA, Räikkönen J, Vucetich LM, Vucetich JA, Peterson RO, Lohmueller KE, et al. Genomic signatures of extensive inbreeding in Isle Royale wolves, a population on the threshold of extinction. Sci Adv. 2019;5: 1–13. doi:10.1101/440511

13. Mukai T, Yamazaki T. The genetic structure of natural populations of *Drosophila melanogaster*, V. coupling-repulsion effect of spontaneous mutant polygenes controlling viability. Genetics. 1968;59: 512–535.

14. Agrawal AF, Whitlock MC. Inferences about the distribution of dominance drawn from yeast gene knockout data. Genetics. 2011;187: 553–566. doi:10.1534/genetics.110.124560

15. García-Dorado A, Caballero A. On the average coefficient of dominance of deleterious spontaneous mutations. Genetics. 2000; 1991–2001. doi:10.1534/genetics.104.027706

16. Peters AD, Halligan DL, Whitlock MC, Keightley PD. Dominance and overdominance of mildly deleterious induced mutations for fitness traits in *Caenorhabditis elegans*. Genetics. 2003;165: 589–599. doi:10.1093/genetics/165.2.589

17. Simmons MJ, Crow JF. Mutations affecting fitness in Drosophila populations. Ann Rev Genet. 1977;11: 49–78.

18. Phadnis N, Fry JD. Widespread correlations between dominance and homozygous effects of mutations: Implications for theories of dominance. Genetics. 2005;171: 385–392. doi:10.1534/genetics.104.039016

19. Wright S. Fisher’s theory of dominance. Am Nat. 1929;63: 274–279.

20. Charlesworth B. Evidence against Fisher’s theory of dominance. Nature. 1979;278: 848–849. doi:10.1038/278848a0

21. Ragsdale AP, Moreau C, Gravel S. Genomic inference using diffusion models and the allele frequency spectrum. Curr Opin Genet Dev. 2018;53: 140–147. doi:10.1016/j.gde.2018.10.001

22. Sawyer SA, Hartl DL. Population genetics of polymorphism and divergence. Genetics. 1992;132: 1161–1176. doi:10.1534/genetics.107.073361

23. Williamson S, Fledel-Alon A, Bustamante CD. Population genetics of polymorphism and divergence for diploid selection models with arbitrary dominance. Genetics. 2004;168: 463–475. doi:10.1534/genetics.103.024745

24. Fuller ZL, Berg JJ, Mostafavi H, Sella G, Przeworski M. Measuring intolerance to mutation in human genetics. Nat Genet. 2019;51: 772–776. doi:10.1038/s41588-019-0383-1

25. Huber CD, Durvasula A, Hancock AM. Gene expression drives the evolution of dominance. Nat Commun. 2018;9: 1–11. doi:10.1038/s41467-018-05281-7

26. Boyko AR, Williamson SH, Indap AR, Degenhardt JD, Hernandez RD, Lohmueller KE, et al. Assessing the evolutionary impact of amino acid mutations in the human genome. PLoS Genet. 2008;4: e1000083. doi:10.1371/journal.pgen.1000083

27. Kim BY, Huber CD, Lohmueller KE. Inference of the distribution of selection coefficients for new nonsynonymous mutations using large samples. Genetics. 2017;206: 345–361. doi:10.1534/genetics.116.197145/-/DC1.1

28. Eyre-Walker A, Woolfit M, Phelps T. The distribution of fitness effects of new deleterious amino acid mutations in humans. Genetics. 2006;173: 891–900. doi:10.1534/genetics.106.057570

29. Tataru P, Mollion M, Glémin S, Bataillon T. Inference of distribution of fitness effects and proportion of adaptive substitutions from polymorphism data. Genetics. 2017;207: 1103– 1119. doi:10.1534/genetics.117.300323/-/DC1.1

30. Veeramah KR, Gutenkunst RN, Woerner AE, Watkins JC, Hammer MF. Evidence for increased levels of positive and negative selection on the X chromosome versus autosomes in humans. Mol Biol Evol. 2014;31: 2267–2282. doi:10.1093/molbev/msu166

31. Auton A, Abecasis GR, Altshuler DM, Durbin RM, Bentley DR, Chakravarti A, et al. A global reference for human genetic variation. Nature. 2015;526: 68–74. doi:10.1038/nature15393

32. Huber CD, Kim BY, Marsden CD, Lohmueller KE. Determining the factors driving selective effects of new nonsynonymous mutations. Proc Natl Acad Sci. 2017;114: 4465–4470. doi:10.1073/pnas.1619508114

33. Haller BC, Messer PW. SLiM 3: Forward genetic simulations beyond the Wright-Fisher model. Mol Biol Evol. 2019;36: 632–637. doi:10.1093/molbev/msy228

34. Haller BC, Messer PW. SLiM 2: Flexible, interactive forward genetic simulations. Mol Biol Evol. 2016;34: 230–240. doi:10.1093/molbev/msw211

35. Haller BC, Messer PW. SLiM 4: Multispecies eco-evolutionary modeling. Am Nat. 2023;201. doi:10.1086/723601

36. Tennessen JA, Bigham AW, O’Connor TD, Fu W, Kenny EE, Gravel S, et al. Evolution and functional impact of rare coding variation from deep sequencing of human exomes. Science (New York, NY). 2012;337: 64–69. doi:10.1126/science.1219240.Evolution

37. Gao Z, Waggoner D, Stephens M, Ober C, Przeworski M. An estimate of the average number of recessive lethal mutations carried by humans. Genetics. 2015;199: 1243–1254. doi:10.1534/genetics.114.173351

38. McCune AR, Fuller RC, Aquilina AA, Dawley RM, Fadool JM, Houle D, et al. A low genomic number of recessive lethals in natural populations of bluefin killifish and zebrafish. Science (80-). 2002;296: 2398–2401. doi:10.1126/science.1071757

39. Kyriazis CC, Robinson JA, Lohmueller KE. Using computational simulations to model deleterious variation and genetic load in natural populations. Am Nat. 2023. 10.1086/726736

40. Morton NE, Crow JF, Muller HJ. An estimate of the mutational damage in man from data on consanguineous marriages. Proc Natl Acad Sci. 1956;42: 855–863. doi:10.1073/pnas.42.11.855

41. Bittles AH, Neel J V. The costs of human inbreeding and their implications for variations at the DNA level. Nat Genet. 1994;8: 117–121. doi:10.1038/ng1094-117

42. Hurst LD, Randerson JP. Dosage, deletions and dominance: Simple models of the evolution of gene expression. J Theor Biol. 2000;205: 641–647. doi:10.1006/jtbi.2000.2095

43. Amorim CEG, Gao Z, Baker Z, Diesel JF, Simons YB, Haque IS, et al. The population genetics of human disease: The case of recessive, lethal mutations. PLoS Genet. 2017;13: 1–23. doi:10.1371/journal.pgen.1006915

44. Wade EE, Kyriazis CC, Cavassim MIA, Lohmueller KE. Quantifying the fraction of new mutations that are recessive lethal. Evolution (N Y). 2023;77: 1539–1549. doi:10.1093/evolut/qpad061

45. Robinson JA, Kyriazis CC, Yuan SC, Lohmueller KE. Deleterious variation in natural populations and implications for conservation genetics. Annu Rev Anim Biosci. 2023;11: 93– 114.

46. Bertorelle G, Raffini F, Bosse M, Bortoluzzi C, Iannucci A, Trucchi E, et al. Genetic load: genomic estimates and applications in non-model animals. Nat Rev Genet. 2022;23: 492–503. doi:10.1038/s41576-022-00448-x

47. Dussex N, Morales HE, Grossen C, Dalén L. Purging and accumulation of genetic load in conservation. Trends Ecol Evol. 2023;38: 961–969. doi:10.1016/j.tree.2023.05.008

48. Dukler N, Mughal MR, Ramani R, Huang Y-F, Siepel A. Extreme purifying selection against point mutations in the human genome. Nat Commun. 2022;13: 1–12. doi:10.1038/s41467-022-31872-6

49. Agarwal I, Fuller ZL, Myers SR, Przeworski M. Relating pathogenic loss-of function mutations in humans to their evolutionary fitness costs. Elife. 2023;12: 1–36. doi:10.7554/elife.83172

50. Agarwal I, Przeworski M. Mutation saturation for fitness effects at human CpG sites. Elife. 2021;10: 1–23. doi:10.7554/eLife.71513

51. Balick DJ, Jordan DM, Sunyaev S, Do R. Overcoming constraints on the detection of recessive selection in human genes from population frequency data. Am J Hum Genet. 2022;109: 33–49. doi:10.1016/j.ajhg.2021.12.001

52. Clark DW, et al. Associations of autozygosity with a broad range of human phenotypes. Nat Commun. 2019;10: 1–17. doi:10.1038/s41467-019-12283-6

53. Narasimhan VM, Hunt KA, Mason D, Baker CL, Karczewski KJ, Barnes MR, et al. Health and population effects of rare gene knockouts in adult humans with related parents. Science (80-). 2016;352: 1312–1316.

54. Szpiech ZA, Xu J, Pemberton TJ, Peng W, Zöllner S, Rosenberg NA, et al. Long runs of homozygosity are enriched for deleterious variation. Am J Hum Genet. 2013;93: 90–102. doi:10.1016/j.ajhg.2013.05.003

55. Fridman H, Yntema HG, Mägi R, Andreson R, Metspalu A, Mezzavila M, et al. The landscape of autosomal-recessive pathogenic variants in European populations reveals phenotype-specific effects. Am J Hum Genet. 2021;108: 608–619. doi:10.1016/j.ajhg.2021.03.004

56. Malawsky DS, Walree E van, Jacobs BM, Heng TH, Huang QQ, Sabir AH, et al. Influence of autozygosity on common disease risk across the phenotypic spectrum. Cell. 2023;186: 4514– 4527. doi:10.1016/j.cell.2023.08.028

57. Johri P, Aquadro CF, Beaumont M, Charlesworth B, Excoffier L, Eyre-Walker A, et al. Recommendations for improving statistical inference in population genomics. PLoS Biol. 2022;20: 1–23. doi:10.1371/journal.pbio.3001669

58. Ragsdale AP. Local fitness and epistatic effects lead to distinct patterns of linkage disequilibrium in protein-coding genes. Genetics. 2022;221. doi:10.1093/genetics/iyac097

59. Garcia JA, Lohmueller KE. Negative linkage disequilibrium between amino acid changing variants reveals interference among deleterious mutations in the human genome. PLoS Genet. 2021;17: 1–25. doi:10.1371/journal.pgen.1009676

60. Barroso G V., Lohmueller KE. Inferring the mode and strength of ongoing selection. Genome Res. 2023;33: 632–643. doi:10.1101/gr.276386.121

61. Gutenkunst RN, Hernandez RD, Williamson SH, Bustamante CD. Inferring the joint demographic history of multiple populations from multidimensional SNP frequency data. PLoS Genet. 2009;5: 1–11. doi:10.1371/journal.pgen.1000695

62. Ségurel L, Wyman MJ, Przeworski M. Determinants of mutation rate variation in the human germline. Annu Rev Genomics Hum Genet. 2014;15: 47–70. doi:10.1146/annurev-genom-031714-125740

63. Jouganous J, Long W, Ragsdale AP, Gravel S. Inferring the joint demographic history of multiple populations: Beyond the diffusion approximation. Genetics. 2017;206: 1549–1567. doi:10.1534/genetics.117.200493

64. Madigan D, Raftery AE. Model selection and accounting for model uncertainty in graphical models using occam’s window. J Am Stat Assoc. 1994;89: 1535–1546. doi:10.1080/01621459.1994.10476894

65. Posada D, Buckley TR. Model selection and model averaging in phylogenetics: Advantages of Akaike information criterion and Bayesian approaches over likelihood ratio tests. Syst Biol. 2004;53: 793–808. doi:10.1080/10635150490522304

66. Keightley PD. Rates and fitness consequences of new mutations in humans. Genetics. 2012;190: 295–304. doi:10.1534/genetics.111.134668

67. Crow JF. Genetic loads and the cost of natural selection. Mathematical Topics in Population Biology.Berlin: Springer; 1970. pp. 128–177.

